# Arid5a mediates an IL-17-dependent pathway that drives autoimmunity but not antifungal host defense

**DOI:** 10.1101/2022.02.14.480386

**Authors:** Tiffany C. Taylor, Yang Li, De-Dong Li, Saikat Majumder, Mandy McGeachy, Partha S. Biswas, Sebastien Gingras, Sarah L. Gaffen

## Abstract

IL-17 contributes to the pathogenesis of certain autoimmune diseases, but conversely is essential for host defense against fungi. Antibody-based biologic drugs that neutralize IL-17 are effective in autoimmunity but can be accompanied by adverse side effects. *Candida albicans* is a commensal fungus that is the primary causative agent of oropharyngeal and disseminated candidiasis. Defects in IL-17 signaling cause susceptibility to candidiasis in mice and humans. A key facet of IL-17 receptor signaling involves RNA binding proteins (RBP), which orchestrate the fate of target mRNA transcripts. In tissue culture models we showed that the RBP AT-rich interacting protein 5a (Arid5a) promotes the stability and/or translation of multiple IL-17-dependent mRNAs. Moreover, during OPC, Arid5a is elevated within the oral mucosa in an IL-17-dependent manner. However, the contribution of Arid5a to IL-17-driven events *in vivo* is poorly defined. Here, we used CRISPR/Cas9 to generate mice lacking Arid5a. *Arid5a*^*-/-*^ mice were fully resistant to experimental autoimmune encephalomyelitis (EAE), an autoimmune setting in which IL-17 signaling drives pathology. Surprisingly, *Arid5a*^*-/-*^ mice were resistant to OPC and systemic candidiasis, similar to immunocompetent WT mice and contrasting with mice defective in IL-17 signaling. Therefore, Arid5a-dependent signals mediate pathology in autoimmunity yet are not required for immunity to candidiasis, indicating that selective targeting of IL-17 signaling pathway components may be a viable strategy for development of therapeutics that spare IL-17-driven host defense.

## Introduction

IL-17 (IL-17A) is a member of a subclass of cytokines that are structurally and functionally distinct from other inflammatory cytokines. Though produced mainly by T lymphocytes (“Type 17” cells, including γδ- and αβ-T cells, ILC3, NKT cells), IL-17 predominantly activates downstream signals in non-hematopoietic cells to protect against mucosal pathogens, notably the commensal fungus *Candida albicans* (1, 2). Conversely, IL-17 and Type 17 cells have been implicated in the pathogenesis of many autoimmune and inflammatory diseases (3), highlighted in the success of anti-cytokine drugs targeting IL-17 or the IL-17 receptor. Though unquestionably these drugs have improved patients’ disease activity and quality of life, their use is associated with adverse side effects including worsening bowel inflammation in patients with inflammatory bowel disease (IBD) and the frequent development of opportunistic fungal infections, especially oral and esophageal candidiasis (4-6).

IL-17 signals through a heterodimeric receptor composed of IL-17RA and IL-17RC, though some alternative configurations of the receptor have recently been described (7, 8). Binding of IL-17 to the IL-17R complex promotes recruitment of the adaptor Act1 (9-11). In turn, Act1 binds to multiple TRAF factors that promote alternative downstream signaling cascades (12). TRAF6 activates several transcription factors, including NF-κB, IκBξ and C/EBP family members. Additionally, Act1 initiates recruitment of TRAF2 and TRAF5, which initiates a signaling cascade resulting in prolonged half-life of many IL-17-dependent mRNA transcripts. The effects of IL-17 signaling on mRNA fate is orchestrated by a complex array of RNA binding proteins (RBPs) (13). RBPs shape the output of inflammatory effector genes by stabilizing or degrading transcripts that encode inflammatory mediators. Some RBPs, including HuR (*Elavl1*) and IMP2 (*Igf2bp2*) act as positive regulators of mRNA expression, resulting in target mRNA stabilization or enhanced translation (14-17). Alternatively, RBPs such as Regnase-1, Splicing factor 2 (SF2) and Roquins negatively regulate mRNA expression, facilitating degradation of unstable transcripts (18-20). These pathways are interlinked because many transcription factors operative in the IL-17 pathway are encoded by RNAs that are subject to post-transcriptional control, for example, IκBξ (*Nfkbiz*) and C/EBPs (*Cebpb, Cebpd*).

AT-rich interaction domain 5A (Arid5a) is another RBP implicated in IL-17 signaling and Th17-driven diseases. Arid5a enhances target mRNA transcript stability and translation, in part by offsetting the destabilizing activity of the endoribonuclease Regnase-1 (14, 21-23). Arid5a controls expression of genes downstream of the Th17/IL-17R axis through control of *Il6* and *Nfkbiz*, as well as Th17 cell responses via *Il6, STAT3, TBX21* (Tbet), and *OX40* transcripts (21, 23, 24). To date, the precise mechanisms by which IL-17 regulates Arid5a function in autoimmunity are not fully understood. Even less is known about Arid5a function in the context of host defense, though Arid5a mRNA is upregulated in an IL-17-dependent manner in oropharyngeal candidiasis (OPC) in the murine oral mucosa (14, 25). Here, we created a novel line of *Arid5a*^*-/-*^ mice to assess the role of Arid5a in autoimmunity versus fungal infections. These data reveal a surprising distinction used between signaling pathways critical in host defense compared to those required in autoimmunity.

## Material and Methods

### Generation of Arid5a^-/-^ mice

The Arid5a knockout allele was fortuitously generated during an attempt to create a conditional deletion with CRISPR/Cas9 (26, 27). Briefly, the targeting strategy was to introduce a LoxP site in intron 2 and a Myc-tag at the C-terminal with another loxP site just after the stop codon. In the process of genotyping potential founders, we identified a mouse carrying a deletion between the two SpyCas9 target sites, with a deletion of 3,359 bp deletion on chromosome 1 between position 36,316,840 and 36,320,199. Fertilized embryos (C57BL/6J, The Jackson Laboratory) produced by natural mating were microinjected in the cytoplasm with a mixture of 0.33 µM EnGen Cas9 protein (New England Biolabs, M0646T), Arid5a-guide2 and Arid5a-guide5 (21.23 ng/µl (∼0.66 µM) and two single stranded oligonucleotides: Arid5a-Myc-HDR and Arid5a-guide5-HDR (0.5 µM, “Ultramers” from IDT). Injected zygotes were cultured overnight, and 2-cell embryos were transferred to pseudopregnant CD1 mice. The sgRNA templates were generated by PCR (26) and used for synthesis of sgRNAs using NEB HiScribe™ T7 Quick High Yield RNA Synthesis Kit (New England Biolabs, E2050S). The sgRNAs purified using MEGAclear Kit (ThermoFisher Scientific). The target sequence of the sgRNAs are: ATAGGCTCTGGCCTACAGTTTGG for Arid5a-guide2 at chr1:36377037-36377060 and GTAAAAGCCAAATGCGCCCCAGG for Arid5a-guide5 at chr1:36373679-36373702 (coordinates from GRCm38/mm10 Assembly).

The sequence of the single-stranded oligonucleotides are: 5’-

tcgaactcagagagttctgcctgcttctcctgagtgctaggattaaaggtgtgtgccaccactgcctgggAATTCATAACTTCGTATAATGTATGCTATACGAAGTTATgcgcatttggcttttaccatacggttgagggactctgaccctgccttccaggaactgagttggataa-3’ for Arid5a-guide5-HDR

5’-TGGCACATGCCACCCGTCACAACCTATGCGGCACCTCACTTCTTCCACCTCAACACCAAACTGGAGCAGAAACTCATCTCAGAAGAGGATCTGTAAGAATTCATAACTTCGTATAATGTATGCTATACGAAGTTATAGAGCCTATCCTGCTATGCTGTGGAGGATTTGATGGGCAGCTGCCGCCATTATCTCAGGCC-3’ for Arid5a-Myc-HDR.

The potential founders were identified by PCR and the sequence of the knockout allele was determined by sequencing. *Arid5a*^*-/-*^ mice or littermates were genotyped with the following primers:

F31-5’-ACCTTCTGGCTACAACAGGC-3’

R31-5’-ACCTACACACTTGCTCCTGC-3’

F51-5’-ATATGGGTGCTAGGCAAGGC-3’

The WT allele is detected with primer F31 and R31 and produces a product of 564 bp. The knockout allele is detected with primer F51 and R31 and produces a product of 595 bp.

### Candidiasis Models

OPC was induced in 7-9-week-old mice of both sexes by sublingual inoculation with *C. albicans* strain CAF2-1-saturated cotton balls for 75 minutes under anesthesia, as described (28, 29). Tongue homogenates were prepared on GentleMACS homogenizers (Miltenyi Biotec) with C-tubes, and CFU was determined by serial dilution plating on Yeast extract-peptone-dextrose (YDP) with ampicillin. Limit of detection is ∼30 CFU/g. For systemic candidiasis, mice were injected i.v. with 2-7 × 10^5^ *C. albicans* strain SC5412 in PBS. Three days post-infection, kidney tissues were homogenized in C-tubes, and CFU was determined as above.

### Experimental autoimmune encephalomyelitis

Mice (males age 6-18 weeks) were immunized subcutaneously in 4 sites on the back with 100 μg MOG peptide 35-55 (Biosynthesis) emulsified with Complete Freund’s Adjuvant (CFA) with *M. tuberculosis* strain H37Ra (DIFCO laboratories) and administered 100 ng pertussis toxin (List Biological Laboratories) i.p. on day 0 and day 2, as described (30). Mice were scored daily as follows: 1. Flaccid tail; 2. Impaired righting reflex and hindlimb weakness; 3. Partial hindlimb paralysis; 4. Complete hindlimb paralysis; 5. Hindlimb paralysis with partial forelimb paralysis; 6. Moribund.

### Flow Cytometry

Tongues were harvested and digested with Collagenase IV (0.7 mg/ml) in Hanks’ balanced salt solution for 30-45 mins at 37°C. Cells were separated by Percoll gradient centrifugation. Abs were from the following sources: anti-CD45 and anti-CD11b (BioLegend), anti-CD4 and anti-F4/80 (Invitrogen), anti-TCRβ, anti-CD8, and anti-CD19 (eBioscience), and anti-Ly6G and anti-Ly6C (BD Biosciences). Dead cells were excluded using Ghost Dye (eBioscience). Data were acquired with an LSRFortessa and analyzed using FlowJo software (TreeStar).

Collected single-cell suspensions from LN were filtered and dead cells were excluded using Ghost Dye (eBioscience). LN were cultured in complete medium (RPMI media containing 10% FCS, supplemented with L-glutamine and antibiotics) with 50 ng/ml PMA and 500 ng/ml ionomycin (Sigma-Aldrich) in the presence of Golgiplug (BD Biosciences) for 4 h, followed by staining with Ghost Dye, CD4 and IL-17 (Biolegend).

### Immune Cell Isolation

Naïve splenocytes and thymocytes were isolated from 6-week old *Arid5a*^*-/-*^ and *Arid5a*^*+/+*^ male mice. Flow cytometry was performed on single cell suspensions.

### Western Blotting

Western blotting was performed as described (31). Abs against Arid5a were from Abcam (ab81149). Blots were imaged with a FluorChem E imager (ProteinSimple, Santa Clara CA).

### Statistics

Data were analyzed on Graphpad Prism. Each symbol represents one mouse unless indicated.

\**P*<0.05, **<0.01, ***<0.001, ****<0.0001. *P* values less than 0.05 were considered significant.

### Study approval

All the experiments were conducted following NIH guidelines under protocols approved by the University of Pittsburgh IACUC.

## Results

### Characterization of Arid5a-deficient mice

To elucidate the role of Arid5a *in vivo*, CRISPR/Cas9 was used to create Arid5a-deficient mice with a deletion in exons 3-7 (**Fig 1A**, see Methods for details). Founders were validated by PCR (**Fig 1B)** and bred to homozygosity. Mice were born at expected Mendelian ratios and exhibited normal fertility and baseline body weights (data not shown). Arid5a is expressed at variable levels across tissues, with high expression in thymus, medium levels in spleen, and very low levels in kidney (**Fig 1C)**. Confirming that mice lacked Arid5a protein expression, tissues from *Arid5a*^*-/-*^ mice displayed undetectable levels of Arid5a compared to *Arid5a*^*+/+*^ mice.

**Figure 1.**
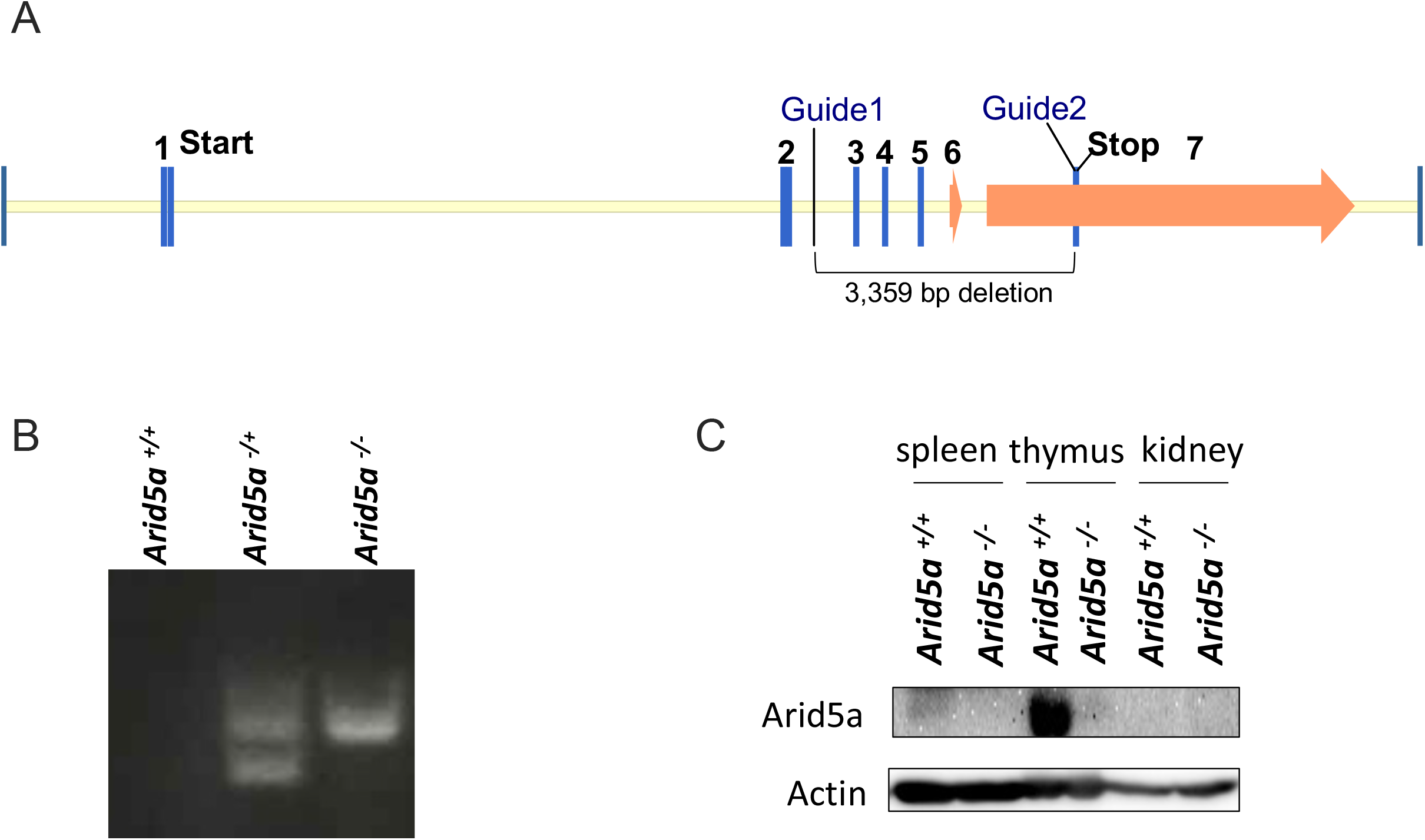
Creation of *Arid5a*^*-/-*^ mice by CRISPR/Cas9. (A) Schematic diagram of Arid5a gene locus and area of deletion. Approximate locations of guide RNA sites are indicated. (B) Genotyping validation of control littermates, heterozygous and homozygous Arid5a-knockout mice. (C) Western blotting of spleen, thymus and kidney tissues from *Arid5a*^*-/-*^ and *Arid5a*^*+/+*^ littermate controls. Top: Arid5a. Bottom: β-actin loading control.

Previous studies using an independently-generated Arid5a knockout mouse demonstrated a role for Arid5a in regulating several immune processes (21-24, 32-34). Therefore, we examined the baseline status of immune compartments in *Arid5a*^*+/+*^ and *Arid5a*^*-/-*^ littermates in spleen (**Fig 2A**) and thymus (**Fig 2B**). As shown, no observable differences between *Arid5a*^*+/+*^ and *Arid5a*^*-/-*^ mice were detected by flow cytometry in thymic or splenic hematopoietic cells, including TCRβ^+^ cells, B cells, neutrophils, monocytes or macrophages. Therefore, the absence of Arid5a does not appear to impact normal development of immune cells.

**Figure 2.**
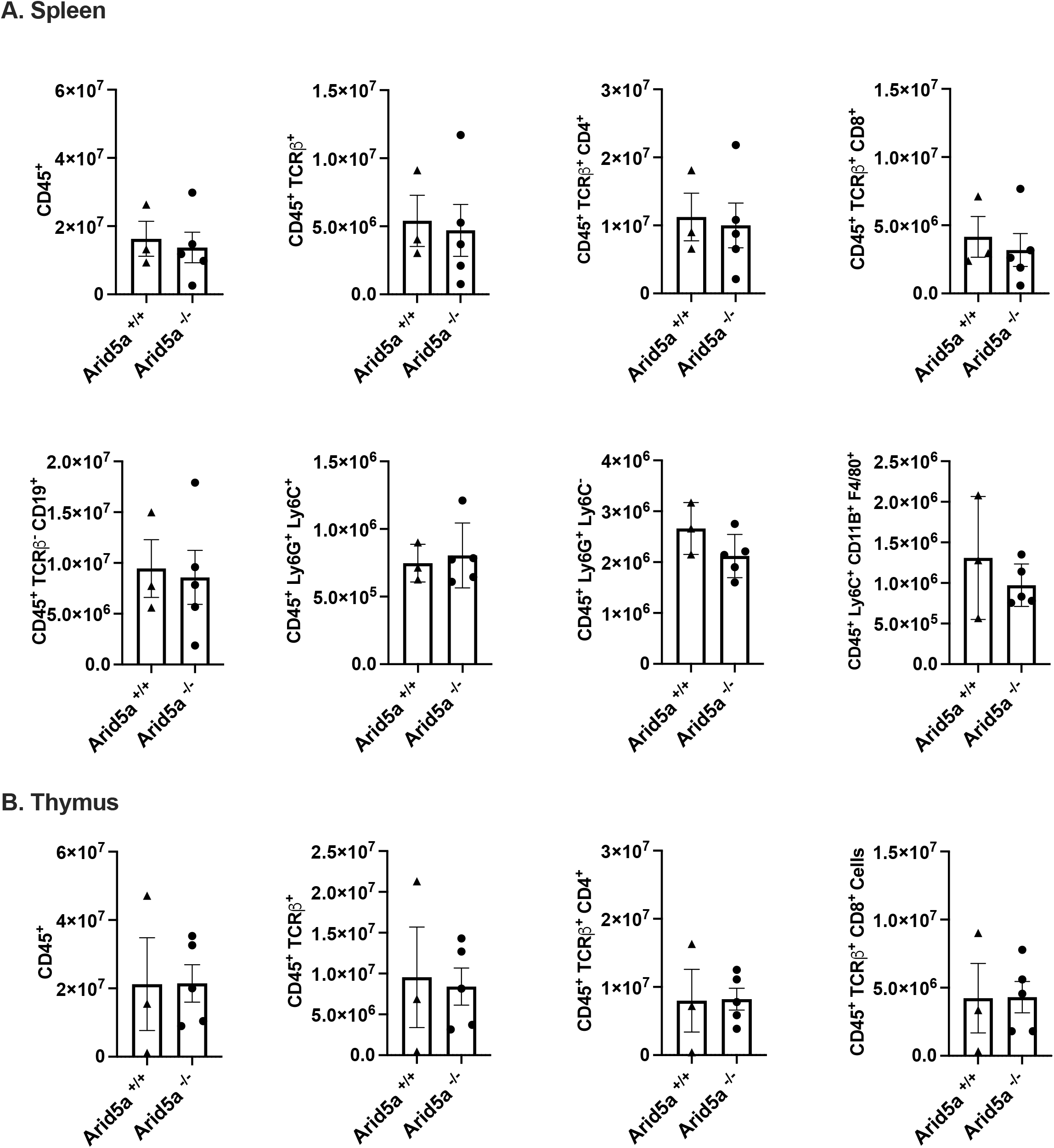
Immunophenotyping of *Arid5a*^*-/-*^ mice. Spleen (A) or thymus (B) from *Arid5a*^*-/-*^ or *Arid5a*^*+/+*^ littermate controls were stained with the indicated cellular markers and analyzed by flow cytometry. Y-axis indicates total cell numbers. Each dot indicates one mouse. Bars indicate mean + SEM. No significant differences between groups were identified by Student’s t-test.

### Arid5a is required for pathogenesis of EAE

Several reports have described a role for Arid5a in driving pathogenesis in mouse models of autoimmunity, with emphasis on control of Th17 cell differentiation (21, 23). Accordingly, we subjected *Arid5a*^*-/-*^ mice and *Arid5a*^*+/+*^ littermate controls to a standard model of experimental autoimmune encephalomyelitis (EAE), a strongly IL-17-dependent model a of multiple sclerosis (16, 35-38). Mice were injected with a myelin oligodendrocyte glycoprotein peptide (MOG 35-55) emulsified with complete Freund’s Adjuvant (CFA). Mice were assessed daily for signs of ascending paralysis on a standard scoring system. *Arid5a*^*+/+*^ control mice developed a typical onset and clinical course of EAE, peaking at 14-16 days post immunization. As expected, mice lacking the IL-17 receptor adaptor, Act1 and *Arid5a*^*-/-*^ were resistant to disease (**Fig 3A**). Concomitant with reduced disease scores, there was marked reduction in EAE incidence in *Arid5a*^*-/-*^ mice (**Fig 3B)**. Th17 cells are pathogenic in EAE (3, 30, 39, 40). To determine if Arid5a affects Th17 cells during EAE, mice were immunized with MOG, and lymph nodes (LN) and spinal cords were harvested on day 12. Cells isolated from LN were stimulated with PMA and stained for CD4 and intracellular IL-17A. There were reduced numbers of CD4^+^ IL-17^+^ cells in *Arid5a*^*-/-*^ compared with *Arid5a*^*+/+*^ mice (**Fig 3C**). Arid5a is known to regulate *Il6* in many cell types (41)(31), and indeed there was a trend to decrease expression in CNS (**Fig 3D**). However, surprisingly, *Stat3* mRNA expression was unchanged (**Fig 3D**) (22). These findings confirm prior work in an independent system that Arid5a is required for development of EAE (41), and also functionally verify that this line of *Arid5a*-knockout mice has the expected phenotype with respect to IL-17-driven autoimmunity.

**Figure 3.**
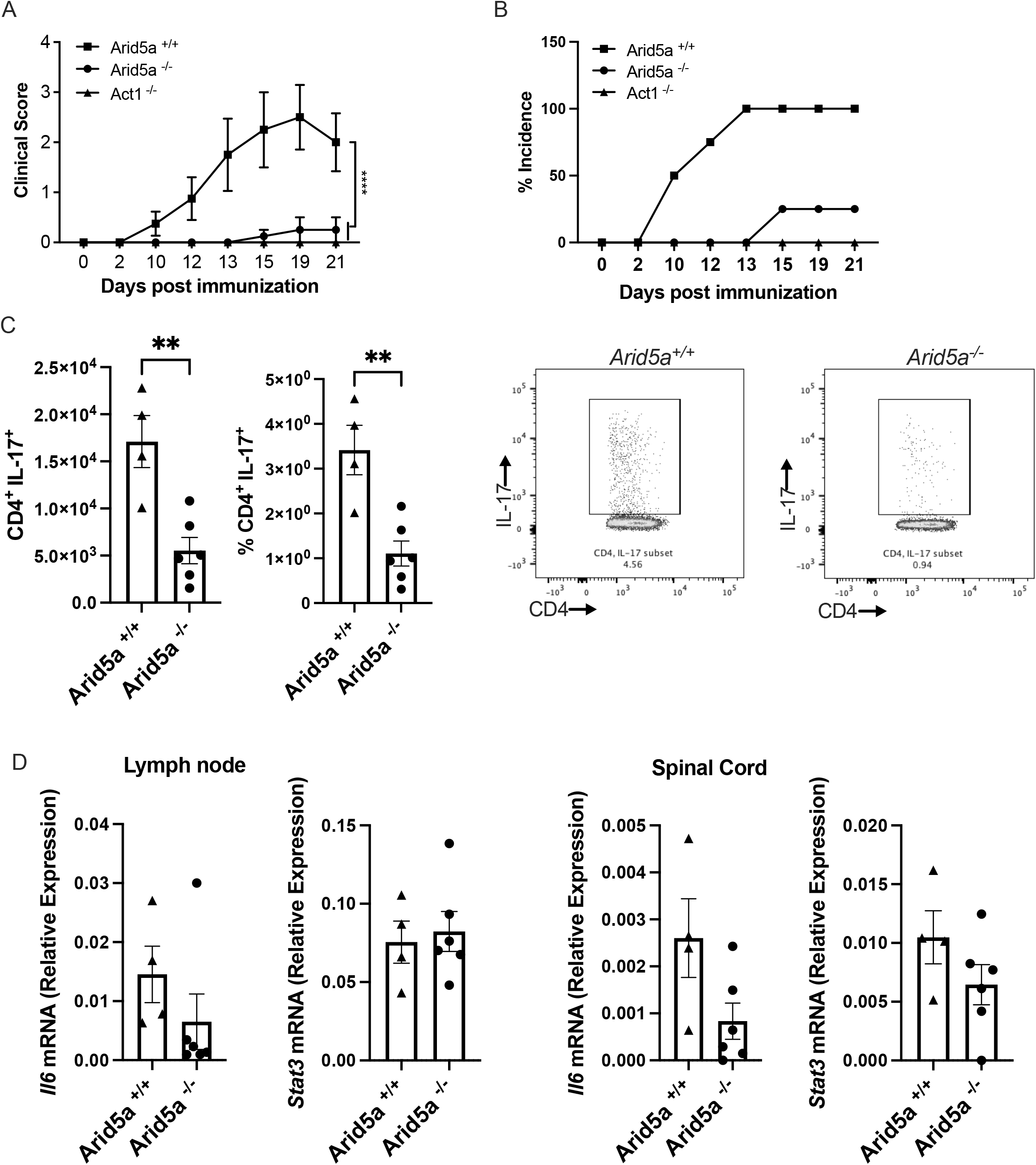
*Arid5a*^*-/-*^ mice are resistant to EAE. The indicated mice (*Arid5a*^+/+^ n=4; *Arid5a*^-/-^ n=4; and *Il17ra*^-/-^ n=3) were subjected to EAE by injection of the MOG peptide. (A) Clinical score was assessed daily by investigators blinded to sample identity. *** P<0.001 by ANOVA with Mann-Whitney U analysis. (B) Percent incidence of EAE in each group is indicated. (C) Lymph node cells harvested on day 12 (*Arid5a*^*+/+*^ n=4 and *Arid5a*^*-/-*^ n=6) were treated with PMA and ionomycin for 4 hours. Cells were stained for CD4 and IL-17A and quantified by flow cytometry. Right: representative FACS plots. ** P<0.003 by Student t-test (D) *Il6* and *Stat3* mRNAs were assessed by qPCR in inguinal lymph nodes and spinal cord at day 12, normalized to *Gapdh*. Data show relative expression ± SEM (*Arid5a*^*+/+*^ n=4; *Arid5a*^*-/-*^ n=6). Data from one experiment (of two) is shown.

### Arid5a is dispensable for immunity to oral and systemic C. albicans infections

IL-17R signaling is well established to drive oral immunity to the commensal fungus *C. albicans*, (2, 42, 43). Moreover, we previously observed that Arid5a mRNA is rapidly upregulated in an IL-17-dependent manner in the oral mucosa of mice during OPC (14). Because Arid5a drives cellular responses to IL-17 through post-transcriptional control of mRNA *in vitro* (14, 44), we predicted that *Arid5a*^*-/-*^ would likely be more susceptible to OPC than WT mice, which clear infection rapidly and do not exhibit overt signs of disease. To test this hypothesis, *Arid5a*^*-/-*^ mice were subjected to OPC by sublingual exposure with a cotton ball soaked with 2 × 10^7^ cells/ml *C. albicans* strain CAF2-1 (29). As a control, *Act1*^*-/-*^ mice (which lack the ability to respond to IL-17 signaling (45, 46)) showed high fungal loads at 5 days after infection (**Fig 4A**) and progressively lost weight throughout the time course of infection (**Fig 4B**). In contrast, *Arid5a*^*+/+*^ and *Arid5a*^*-/-*^ mice cleared the infection by day 5 and returned to normal body weights (**Fig 4A, B**).

**Figure 4.**
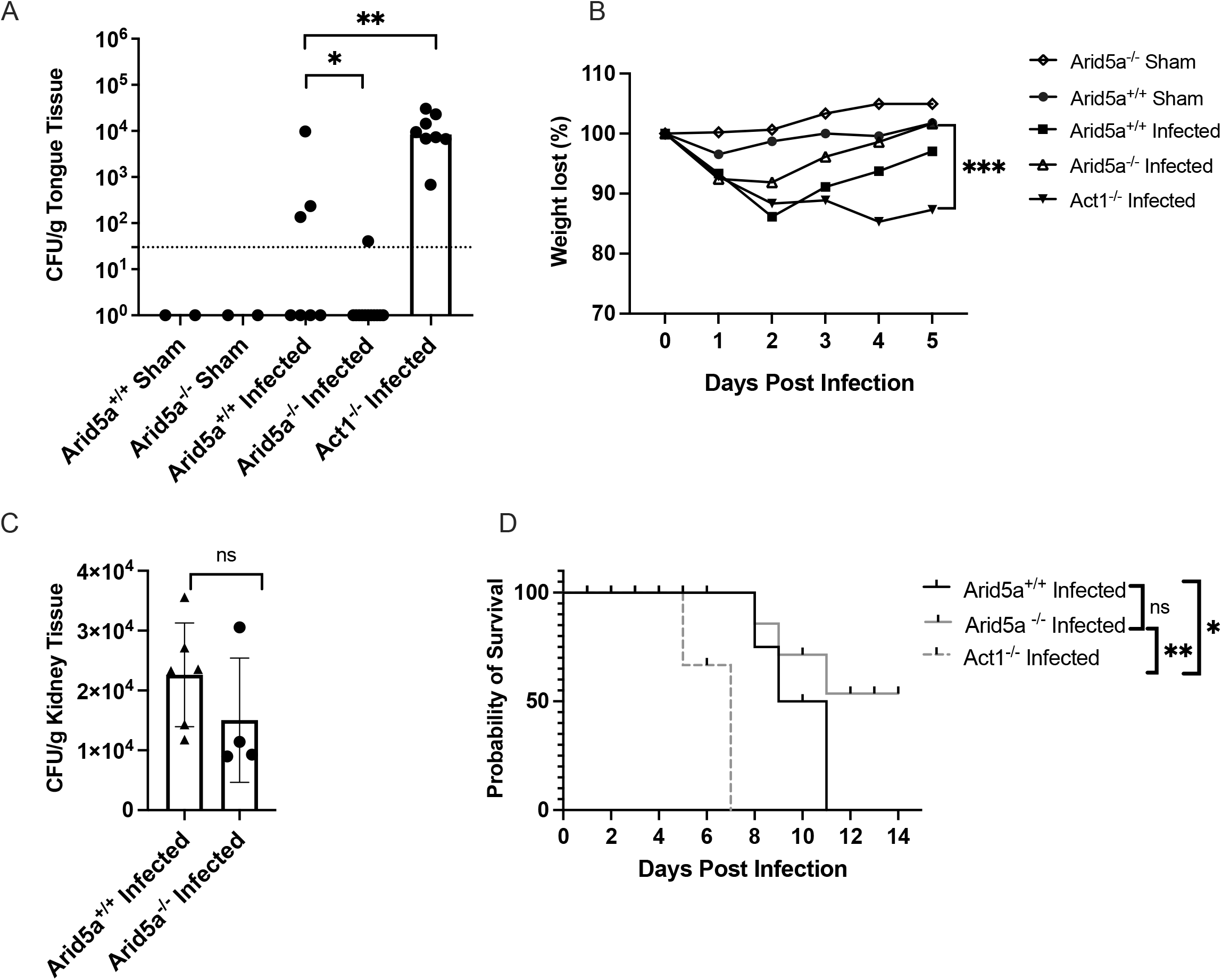
*Arid5a*^*-/-*^ mice show no alterations in susceptibility to mucosal or systemic candidiasis. (A) The indicated mice were infected orally with PBS (Sham) or *C. albicans* strain CAF2-1 for 75 min. After 5 days, tongues were assessed for CFU by plating. Dashed line = limit of detection. Each dot indicates one mouse. Bars indicate geometric mean. *P<0.05, ****P<.0001 by ANOVA with Kruskal-Wallis test (B) The mice in panel A were weighed daily and percent weight loss relative to day 0 is indicated. (C) The indicated mice (*Arid5a*^*+/+*^ n=6 and *Arid5a*^*-/-*^n=4) were infected i.v. with *C. albicans* strain SC5314 and on day 3 fungal loads in kidney homogenates were assessed by plating. Experiment was performed once. (D) The indicated mice (*Arid5a*^*+/+*^ n=5, *Arid5a*^*-/-*^ n=6 and Act1^*-/-*^ *n=4*) were infected i.v. with *C. albicans* strain SC5314 and survival was monitored up to 15 days. Data were analyzed by log-rank (Mantel-Cox) test. *P<0.05 **P<0.001. ns, not significant. Experiment was performed twice.

Disseminated candidiasis is a serious nosocomial infection associated with high rates of morbidity and mortality in humans (2). In mouse models, IL-17R signaling mediates immunity to candidiasis (47, 48). Because Arid5a controls genes that drive Th17 differentiation as well as genes downstream of the IL-17 receptor (12, 44), we postulated that Arid5a deficiency would render mice susceptible to disseminated candidiasis. Therefore, we subjected *Arid5a*^*-/-*^ mice to disseminated candidiasis by intravenous inoculation of 2 × 10^5^ *C. albicans* yeast cells (strain SC5314). *Arid5a*^*-/-*^ mice had similar fungal loads as *Arid5a*^*+/+*^ mice (**Fig 4C**). Moreover, while *Act1*^*-/-*^ mice succumbed to disease by day 7, most *Arid5a*^*+/+*^ and *Arid5a*^*-/-*^ mice survived beyond day 10 and did not show statistically different survival patterns, indicating that Arid5a is not required for immunity to systemic *C. albicans* infections (**Fig 4D**). Collectively these data indicate that Arid5a is important for autoimmunity but does not play an important role in the response to *C. albicans*.

## Discussion

Arid5a was originally described for its ability to interact with AT-rich DNA elements in DNA, modulating cellular proliferation, differentiation, and development (49-51). Arid5a belongs to the ARID family that consists of 15 proteins, which can be divided into seven sub-groups based on the degree of similarity within the ARID domain. The ARID domain is a helix-turn-helix motif essential for binding to DNA elements (49, 50) and can also interact with cis-acting elements located in the 3’ UTR of certain inflammatory RNAs, such as *Il6, Tbx21, Stat3, Ox40* (14, 21-24, 52). Binding of Arid5a to these transcripts promotes their longevity and/or facilitates their translation, though the underlying mechanisms are not well understood. Additionally, Arid5a offsets the action of a potent endoribonuclease, Regnase-1, which degrades target transcripts and thereby serves as a negative regulator of inflammation (21). Arid5a acts in hematopoietic cells (macrophages, T cells) and non-hematopoietic cells (fibroblasts, epithelial cells) (12, 44). Thus, Arid5a is a newly recognized RBP that is an important post-transcriptional modulator in multiple immune pathways.

The *Arid5a*^*-/-*^ mice described here are not identical to the original published *Arid5a*^*-/-*^ mouse strain (21). The latter lacks exons 1-3 and a major part of the DNA binding domain was replaced by a floxed Neo-cassette. In the *Arid5a*^*-/-*^ mice described here, LoxP sites were introduced into intron 2 and immediately following the stop codon, creating a 3,359 bp deletion. Despite these differences, both *Arid5a*^*-/-*^ lines are resistant to EAE (41), lending confidence that Arid5a plays an important role in driving Th17/IL-17-driven autoimmune inflammation.

IL-17 mediates effects through upregulation of target RNA accumulation, which can occur transcriptionally but also post-transcriptionally (7). Even so, the data presented here indicate that *Arid5a*^*-/-*^ mice are, unexpectedly, resistant to oral and systemic candidiasis even though IL-17R signals are essential for immunity to this condition (47, 53, 54) and Arid5a functions downstream IL-17 to upregulate many IL-17 target genes (25, 31). Arid5a has been shown to act in T cells to stabilize genes such as *Il6, STAT3, Nfkbiz, OX40, Tbet* that drive Th17 differentiation and IL-17 production (22, 44). One explanation to reconcile these observations may be that the early responses to *C. albicans* in mice are innate in nature, as this fungus is not a commensal microbe in mice (55). Rather than conventional antigen-specific Th17 cells, IL-17 in these naïve settings comes primarily from a combination of γδ-T cells and innate-acting CD4^+^αβ-T cells; additionally, adaptive T cell memory responses can be established in mice that appear to reflect human T cells responses (56-62). In line with this, *Il6*^*-/-*^ and *Ccr6*^-/-^ mice are resistant to OPC, and Th1 cells (IL-12) are dispensable for protection against oral *C. albicans* (28, 56, 63). Thus, the genes regulated by Arid5a might not be important for protection against *Candida albicans* oral infections.

Anti-IL-17 therapy is not associated with risks to systemic candidiasis, likely because the major mechanisms of antifungal control at cutaneous/mucosal surfaces is through neutrophil recruitment and β-defensin production whereas circulating neutrophils are less IL-17-dependent (4, 54, 64, 65). In contrast, during systemic candidiasis IFN-γ, IL-12 as well as IL-17 signaling contribute to effective immunity against *C. albicans* (66-68). Previous studies showed that Arid5a regulates expression of IFN-γ in Th1 cells via control of the master Th1 transcription factor Tbet (24). Given this, we were surprised to find that *Arid5a*^*-/-*^ mice showed the same susceptibility to systemic candidiasis as their wild type littermate counterparts. It is possible that other mRNA stabilizing factors either in T cells or in IL-17-responsive cells [likely, renal tubular epithelial cells (48), though NK cells have been reported (69)] can compensate for the loss of Arid5a during *C. albicans* infection, but more work will be needed to dissect this phenomenon.

The use of biologics targeting IL-17 or IL-17 receptor was a major advance in the treatment of psoriasis, psoriatic arthritis, and ankylosing spondylitis (70). Unexpectedly, IL-17 blockade failed in Crohn’s disease, which has been attributed to a vital tissue protective role of IL-17 in the intestine epithelium (5, 71-73). Intriguingly, IL-17 has been linked to controlling social behavior in mice (74), and blockade of IL-17RA was linked to a small risk of suicide in humans (75), though this is controversial (76). Thus, an ideal therapeutic approach to blocking IL-17 would inhibit its pathogenic effects while sparing host-defensive pathways. Based on these and other data, Arid5a may be an attractive candidate for selective blockade in IL-17-driven diseases, though achieving this is not trivial. Interestingly, recent advances in approaches to understand and target RNA binding proteins suggests potential ways to disrupt Arid5a (or other RBP) interactions with specific target transcripts (77). For example, an elegant study in the IL-17 system showed that aptamers that recognize the binding element of Act1 can interfere with its mRNA stabilizing capacity, and are effective at reducing inflammation *in vivo* (78). Indeed, the development and potential uses of RNA therapeutics is becoming increasingly appreciated (79). Arid5a was reported to be a target of the antipsychotic drug chlorpromazine (22), though this remains to be confirmed. Targeting individual pathway components such as Arid5a potentially allows for a more targeted approach to treat IL-17-mediated inflammatory/autoimmune diseases while preserving the protective role of IL-17 in host defense.

## Acknowledgements

This work was supported by NIH grants to SLG (AI144436, AI147383, DE022550). PSB was supported by AI142354 and AI159058, and MJM by AI148356. TCT was supported by T32-AI089443. We thank B.M. Coleman, N. Amatya and R. Bechara for valuable input.

## Author Contributions

Conceptualization: SLG

Methodology, SG, SLG

Investigation: TCT, YL, DL, SM, PSB, SG

Writing – original draft, TCT, SLG

writing-review and editing, PSB, SG, SLG, MJM

Funding Acquisition, SLG, PSB, MJM

Supervision, SLG, PSB, MJM

